# Blood-based Nano-QuIC: Inhibitor-resistant Detection of Seeding Activity in Patients Diagnosed with Parkinson’s Disease

**DOI:** 10.1101/2023.08.09.552630

**Authors:** Peter R. Christenson, Hyeonjeong Jeong, Manci Li, Hyerim Ahn, Ann M. Schmeichel, Pinaki Misra, Danni Li, Rodolfo Savica, Phillip A. Low, Wolfgang Singer, Peter A. Larsen, Hye Yoon Park, Sang-Hyun Oh

## Abstract

A hallmark of α-synucleinopathies (e.g. Parkinson’s disease) is the misfolding and aggregation of α-synuclein in tissues and biological fluids. Protein amplification assays like real-time quaking-induced conversion (RT-QuIC) are sensitive yet currently limited to semi-invasive sample types such as cerebrospinal fluid because more accessible samples, such as blood, contain inhibitors. Here, we show that Nanoparticle-enhanced Quaking-induced Conversion (Nano-QuIC) can double the speed of reactions spiked with misfolded α-synuclein while increasing sensitivity 100-fold in human plasma. Nano-QuIC detected spike concentrations down to 90 pg/ml in lysed whole blood, while reactions without nanoparticles (RT-QuIC) failed to have any detection due to the presence of strong inhibitors. Moreover, Nano-QuIC showed increased seeding activity in plasma samples from Parkinson’s patients (n=4) versus healthy controls (n=4). This sets the groundwork for the noninvasive diagnostic use of Nano-QuIC, potentially enabling early disease detection and management through blood-based testing.

**TOC graphic:** 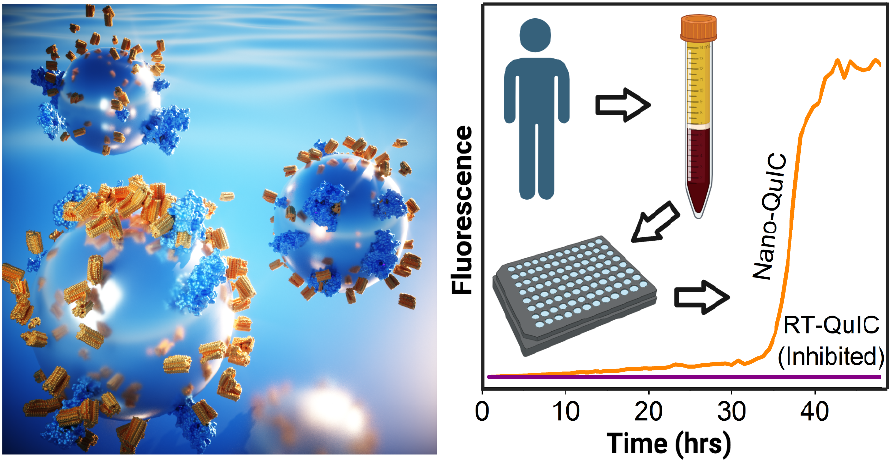

---

The misfolding of native α-synuclein (α-syn) protein is a hallmark of a group of neurodegenerative disorders, including Parkinson’s disease (PD), dementia with Lewy bodies (DLB), and multiple system atrophy (MSA)^1–4^, collectively referred to as α-synucleinopathies. In these disorders, native α-syn misfolds and aggregates into degradation-resistant amyloid deposits that are rich in β-sheet structures. The misfolded α-syn acts as a structural template, triggering other functional α-syn to misfold, thus propagating amyloid deposits, thereby causing subsequent cellular damage. While the specific roles and mechanisms of α-syn misfolding in the pathogenesis of these diseases remain elusive, there is general consensus on the potential for misfolded α-syn to be leveraged as a biomarker for diagnosis, disease progression monitoring, and therapeutic response assessment.^3,5–7^ As such, the early and prodromal detection of misfolded α-syn for clinical applications has become a compelling area of research for α-synucleinopathies.^8–11^

Tremendous progress has been made in the quest for enhanced detection of misfolded α-syn. Currently, seeded amplification assays (SAAs), such as protein misfolding cyclic amplification (PMCA) and real-time quaking-induced conversion (RT-QuIC), outperform traditional immuno-based assays such as ELISA^12^. Additionally, progress has been made utilizing microfluidics for misfolded protein detection.^13^ SAAs leverage the replicating nature of misfolded proteins incubated and agitated in solutions containing an excess of recombinant α-syn. If misfolded α-syn is present in the sample, it acts as a template seed that causes the recombinant α-syn to misfold, thus greatly increasing the number of fibrils in solution.^14^ By including amyloid-binding fluorescent dyes such as thioflavin T (ThT), the amount of misfolded α-syn in the reaction can be monitored in real time. SAA’s like RT-QuIC have demonstrated success for detecting misfolding proteins, including α-syn, in a wide range of neurodegenerative diseases involving protein misfolding.^14,15^ As for α-synucleinopathies, SAAs have detected the presence of pathological α-syn in several sample types including olfactory mucosa, saliva, skin, and CSF at different stages of the disease; it can even differentiate among various α-synucleinopathies using CSF.^12,16–20^

While considerable progress has been made in the clinical detection of misfolded α-syn in the CSF^6,21,22^, the associated collection procedure of CSF, is a painful and pseudo-invasive process that can result in bleeding into the spinal canal, extreme headaches, infections as well as a host of other symptoms. In part due to these issues, CSF is not collected during routine check-ups, limiting its utility for early-stage detection of an α-synucleinopathy.

Given these challenges, the prospect of using non-invasive, readily accessible peripheral fluids such as saliva, urine, or blood^23,24^, for early detection of α-synucleinopathies is highly attractive. Specifically, blood stands out as a particularly desirable target due to its frequent use in routine medical testing and the wealth of biological information it carries. Therefore, our goal in this study is to explore techniques that enable the detection of misfolded α-syn in blood samples.

Among existing assays, SAAs stand as the most promising technique in this regard, but it does face significant challenges, particularly when applied to complex peripheral fluids, such as saliva, urine, and blood.^12^ These fluids contain various components, known as inhibitors, which interfere with the seeding activity in RT-QuIC and PMCA, thus hindering their direct use for the assay.^25–28^ Such inhibitors have been most comprehensively documented and, at times successfully overcome, in studies of Chronic Wasting Disease (CWD).^29–33^ Similarly for α-synucleinopathies, complex and costly steps, such as protein extraction from exosomes and immunoprecipitations (IP)^8,34,35^, were needed prior to amplification using agitated incubation or RT-QuIC, to circumvent the inhibitors present in blood samples.

In short, the quest to make RT-QuIC a clinically viable and translationally successful technology hinges on the ability to overcome the effects of these inhibitors in blood and other biological fluids and expediting reaction times. This focus remains imperative irrespective of the specificity among different α-syn isoforms and diseases.

Nanoparticles have been used to advance diagnostics in a number of fields.^36–38^ Recently, we developed nanoparticle-enhanced quaking-induced conversion (Nano-QuIC) assay, which combines nanotechnology with traditional RT-QuIC to accelerate reaction rates, improve the sensitivity of detection, and crucially, overcome inhibitory components found in complex tissue samples.^37^ Given this advancement, we sought to examine the utility of Nano-QuIC for the detection of misfolded α-syn in blood samples. Blood, owing to its non-invasive collection method and abundant biological information it provides, stands as an ideal candidate for early disease diagnostics.

Here we study the effects of size and concentration of silica nanoparticles (siNPs) in the Nano-QuIC assay for blood samples. We performed tests with siNP diameters ranging from 20-100 nm with concentrations ranging from 0.1-2.5 mg/ml. We then examined the effects of pH on the α-syn-siNP interaction in QuIC assays. Since we focus on overcoming inhibitor effects, and multiple strains and diseases create variations that complicate interpretations, we applied Nano-QuIC to detect misfolded α-syn in complex human plasma and bovine blood spiked with synthetic α-syn seeds (Fig. 1a-c). We show that Nano-QuIC can readily surmount inhibitors and detect 900 pg/ml and 90 pg/ml of spiked α-syn seeds in human plasma and whole lysed bovine blood, respectively. Finally we detected increased seeding activity in plasma from patients diagnosed with PD compared to healthy controls. These proof-of-concept results demonstrate Nano-QuIC’s potential to detect low levels of misfolded α-syn in plasma, an easily accessible sample type.

**Figure 1:**
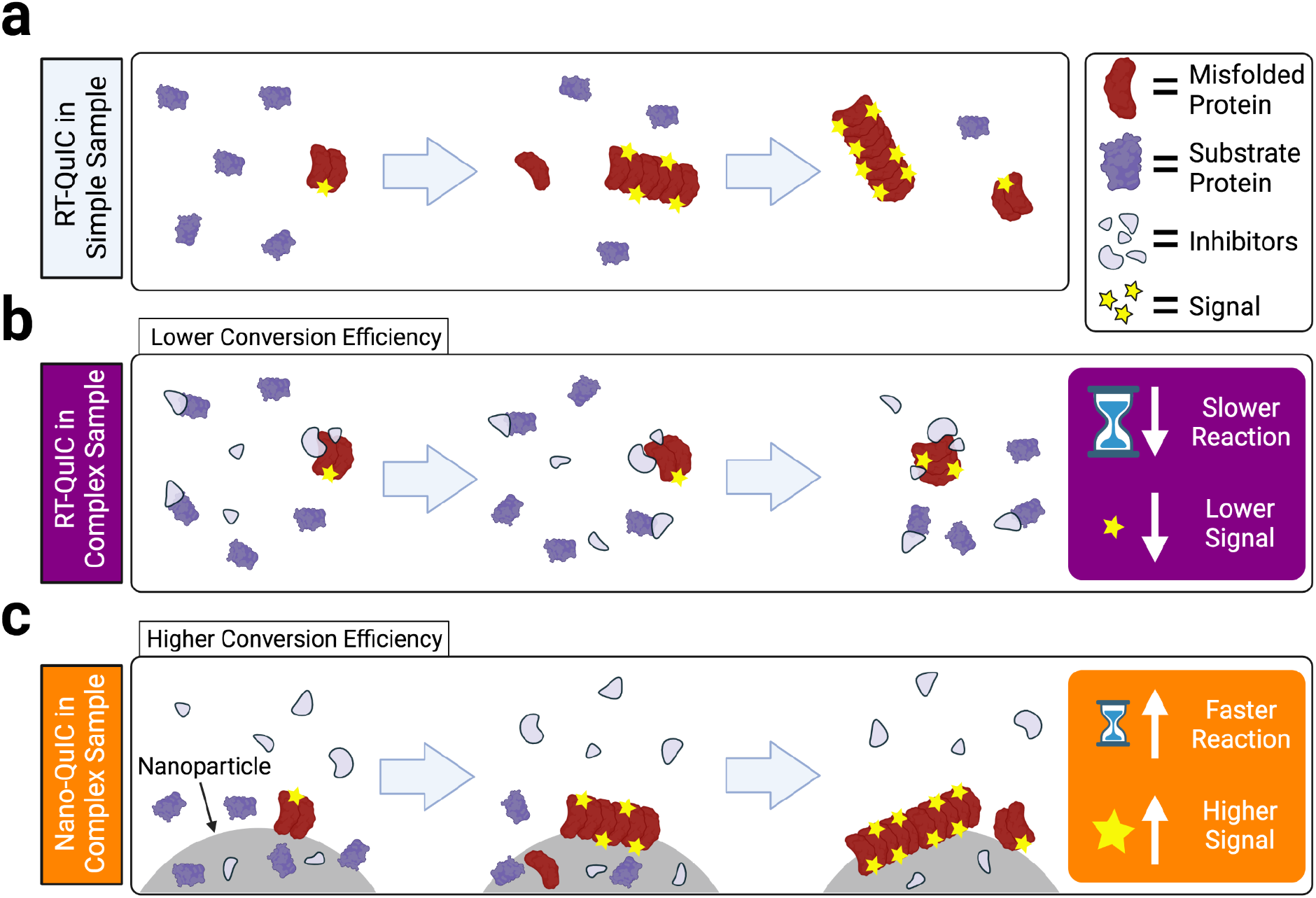
Overview of RT-QuIC vs. Nano-QuIC. (a) Schematic view of RT-QuIC with simple solutions. Misfolded protein catalyzes the substrate protein to misfold, triggering a chain reaction of misfolding events. (b) Possible molecular view of RT-QuIC with complex solution (e. g. whole blood or plasma) inhibiting fibril growth and detection. (c) Proposed molecular view of Nano-QuIC overcoming inhibitors in complex solution. Substrate protein adsorbs onto and/or are concentrated near the surface of the silica nanoparticle, where it is destabilized allowing for the misfolded protein to act as a template for further misfolding events.

## Results

### Diameter and Concentration Effects of Silica NPs (siNPs)

To examine the effect of the diameter and concentration of the siNPs, we performed Nano-QuIC across various diameters (20, 50, and 100 nm) and concentrations (0.1, 0.5, and 2.5 mg/mL) for each diameter. Nano-QuIC was run for 48 hrs at 42 °C and the rate of amyloid formation (RAF, the inverse in seconds of the time it takes for the fluorescence to exponentially rise.) was measured. Compared with the RAF measured without siNPs, RAF with the siNPs were all improved regardless of the diameters and concentrations (Fig. 2a).

**Figure 2:**
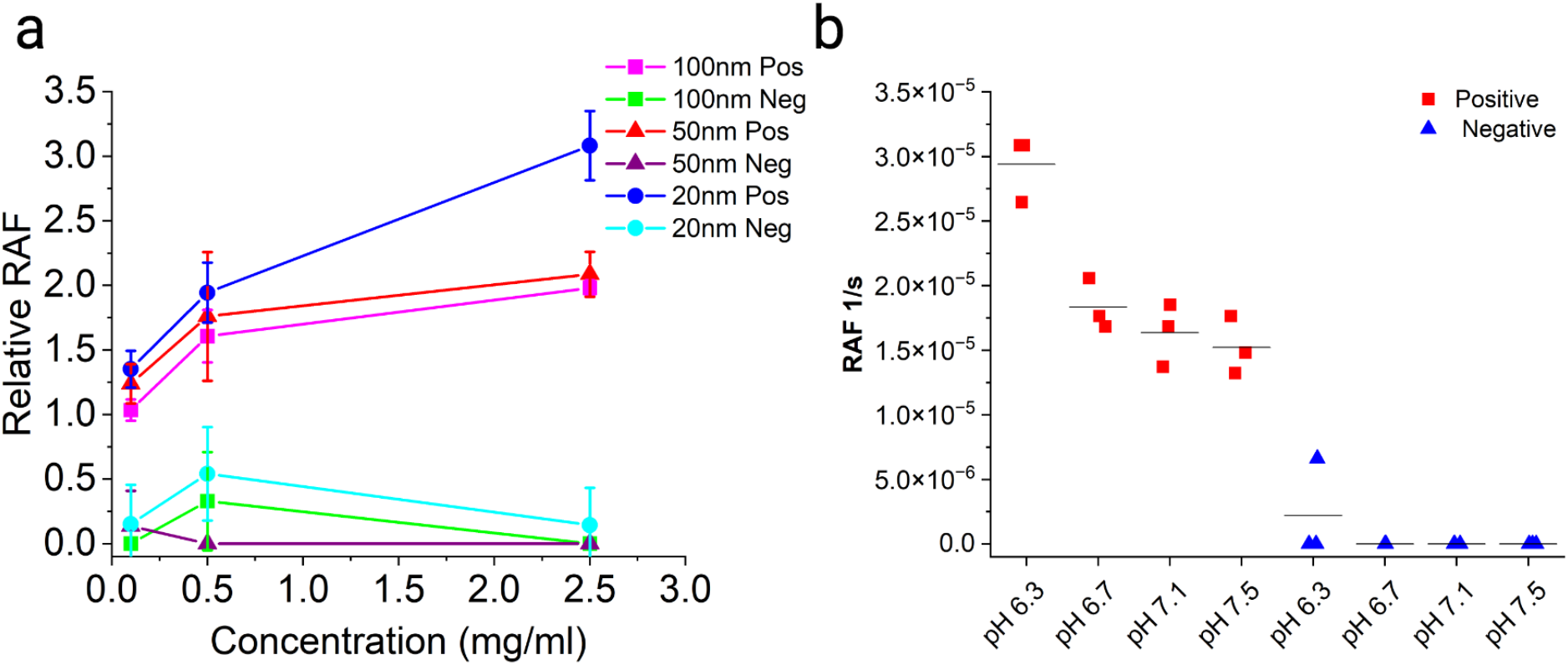
(a) SiNP concentration and diameter sweeps for α-syn Nano-QuIC. Relative RAF is the ratio of the rate of amyloid formation for wells containing siNPs divided by wells without siNPs. “Pos” indicates wells with synthetic misfolded α-syn. “Neg” indicates wells without synthetic misfolded α-syn. All error bars indicate standard deviation. (b) Effects of pH on the RAF of α-syn Nano-QuIC reactions. The speed of the reaction increases as the pH drops.

As the diameter of siNPs became smaller, the RAF value became larger. However, in the RAF of 20 nm siNPs’, false seeding was observed in some of the negative controls, which would lead to lower accuracy of misfolded α-syn detection. On the other hand, in the same diameter of siNPs, as concentration increases, the RAF also improved (Fig. 2a). As a result, 50 nm siNPs, with 2.5 mg/mL was deemed best because it gave significant improvement to the RAF while showing no false seeding.

### Investigating pH Effects

The pH of reactions has been shown to play an important role in seeded amplification assays.^14,39^ There is variation among amplifications assays in the pH used ^12,27,39^. To determine the effects of pH on Nano-QuIC, we performed Nano-QuIC across a pH range of 6.3 to 7.5 for 50 nm siNPs at a concentration of 2.5 mg/ml (Fig. 2b). This range was selected within the buffering capacity of phosphate buffers. We then measured the RAFs and found that as the pH of the master mix solution became lower, the RAF values were increased (Fig. 2b), with the fastest detection time (9.5 hrs) measured at pH 6.3. Notably, in the negative control wells that were not seeded with misfolded α-syn, no seeding activity was detected before the 42 hr mark and only a single technical replicate after that showed low seeding activity. As a result, pH 6.3 for the master mix was considered optimal for detecting misfolded α-syn.

### Detection of misfolded a-syn in human plasma

In previous work, we showed that Nano-QuIC greatly reduces the effect of inhibitors in the lymph tissue of wild white-tailed deer infected with CWD.^37^ In this study, we extended our focus to human plasma samples. To create synthetic misfolded α-syn spikes, RT-QuIC reactions with concentrations of 0.09 mg/ml α-syn were shaken till spontaneous aggregation occurred. These spontaneous seeds were then aliquoted and stored for future use.

Misfolded α-syn spikes in plasma at concentration of 9 μg/ml were serially diluted 10-fold in human plasma down to 90 pg/ml. Each of the 10-fold dilution series was diluted 100-fold in SDS PBS and added to both RT-QuIC and Nano-QuIC reactions. No other extractions or sample preparation steps were performed. We found that Nano-QuIC could detect the misfolded α-syn down 900 pg/ml, whereas RT-QuIC only detected seeding down to 90 ng/ml (Fig. 3a-d). Importantly, there was no false seeding shown in the negative control (Fig. 3a & S1a). Additionally, for all wells showing seeding activity, Nano-QuIC detection time was on average 2X faster than reactions with no SiNPs of the same spike dilution (Fig. 3a-d). Although ∼20 hours average detection time for Nano-QuIC still seems lengthy, this represents a significant gain in throughput since the average reaction time is cut by half from ∼40 hours (for RT-QuIC) to 20 hours (for Nano-QuIC).

**Figure 3:**
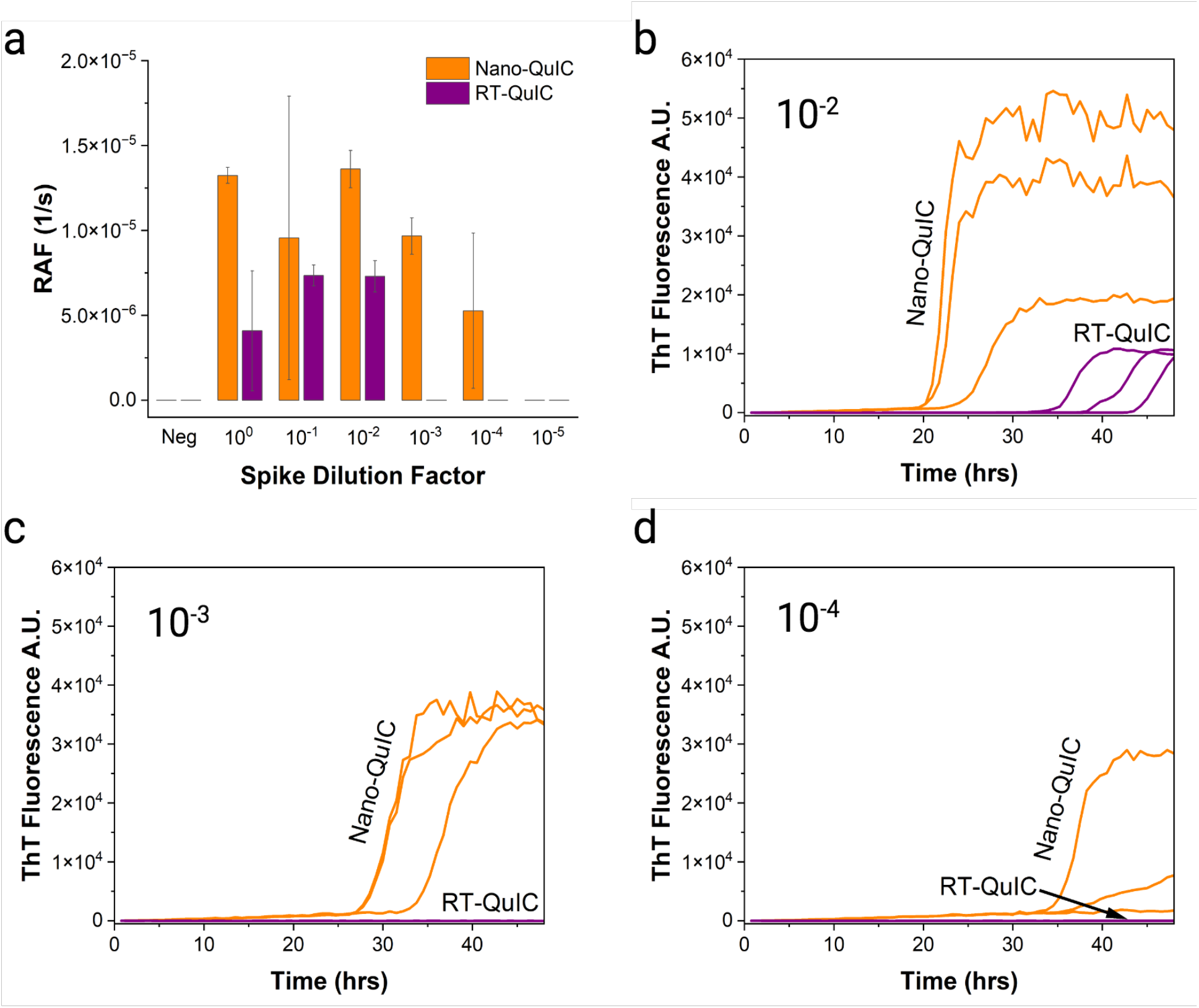
(a) Dilution of misfolded α-syn spike in human plasma from 9 μg/ml to 90 pg/ml. Nano-QuIC overcomes inhibitors to allow a 100-fold improvement in sensitivity. (b-d) ThT fluorescence curves for human plasma samples spiked to 90ng/ml (dilution 10^−2^), 9ng/ml (dilution 10^−3^), and 900pg/ml (dilution 10^−4^) misfolded α-syn.

### Detection of misfolded α-syn in bovine whole blood

Our experiments using human plasma showed proof-of-concept results for using Nano-QuIC to overcome blood-borne inhibitors to detect misfolded α-syn even in complex biological samples. Historically, blood has been a challenging sample type for QuIC-like reactions due to the presence of various inhibitors.^25,40,41^ To further validate the robustness of Nano-QuIC and its ability to negate these inhibitory effects, we expanded our testing scope to include misfolded α-syn spike into lysed whole bovine blood. We utilized bovine whole blood for two reasons. First it presents an even higher level of complexity than human plasma, showcasing the ability of Nano-QuIC to function effectively in extremely challenging biological matrices. Second, the ease of acquiring a large volume of bovine blood allowed us to run a series of Nano-QuIC experiments to develop protocols.

Misfolded α-syn was spiked into bovine blood to a concentration of 9 μg/ml. This sample was then serially diluted 10-fold in bovine blood down to 90 pg/ml. Each of the 10-fold dilution series was further diluted 100-fold in SDS PBS and added to both RT-QuIC and Nano-QuIC reactions. No other extractions or sample preparation steps were performed. Our results demonstrate that Nano-QuIC was able to detect the misfolded α-syn down to a concentration of 90 pg/ml. In stark contrast, reactions with no SiNPs failed to detect any presence of misfolded α-syn due to the presence of inhibitors (Fig. 4a-b). Importantly, our negative control showed no seeding, underscoring the specificity of our Nano-QuIC assay (Fig. 4a & S1b).

**Figure 4:**
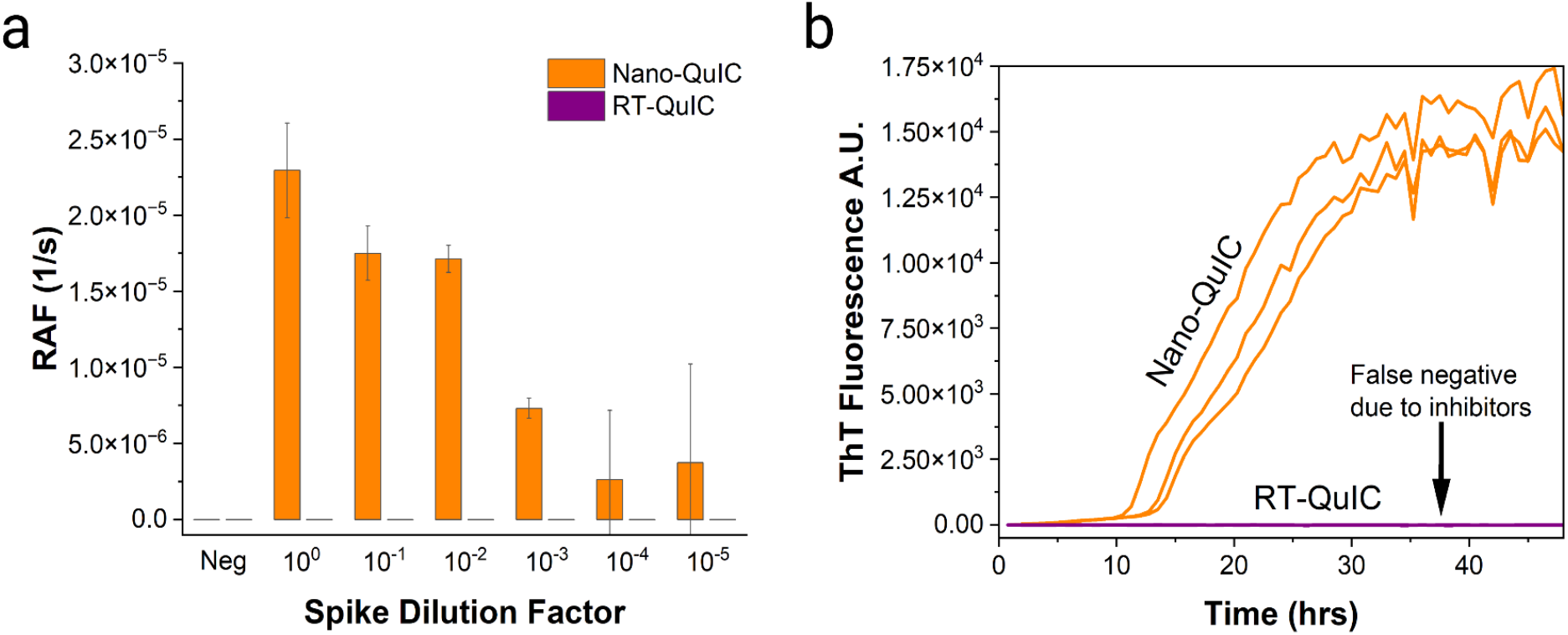
(a) Dilution of misfolded α-syn spike in bovine whole blood from 9 μg/ml to 90 pg/ml. While RT-QuIC shows no detection, Nano-QuIC shows detection down to 90 pg/ml (b) ThT fluorescence curves for bovine blood samples spiked to 9 μg/ml (dilution 10^0^) misfolded α-syn.

### Detection seeding activity from human Parkinson’s patient plasma samples

To demonstrate initial proof of concept with actual patient synucleinopathy samples, Nano-QuIC was performed on individual patient plasma samples generously provided to us by the Mayo Clinic. Four patients with idiopathic PD (3 male, 1 female), median age 68.3 years (range 63.4-74.7) (Fig. 5a-d) and four control subjects without underlying neurologic disease (2 male, 2 female), median age 69.5 years (range 60.1-72.1) (Fig. 5e-h) were included. Samples were collected between May and September 2019. At the time of blood collection, PD patients were between 5 and 10 years from symptom onset and classified as stage Hoehn and Yahr I (n=1) and II (n=3). Patients had between 2.5 and 4.8 years of neurologic follow-up after sample collection and their clinical course remained consistent with the diagnosis of PD.

**Figure 5:**
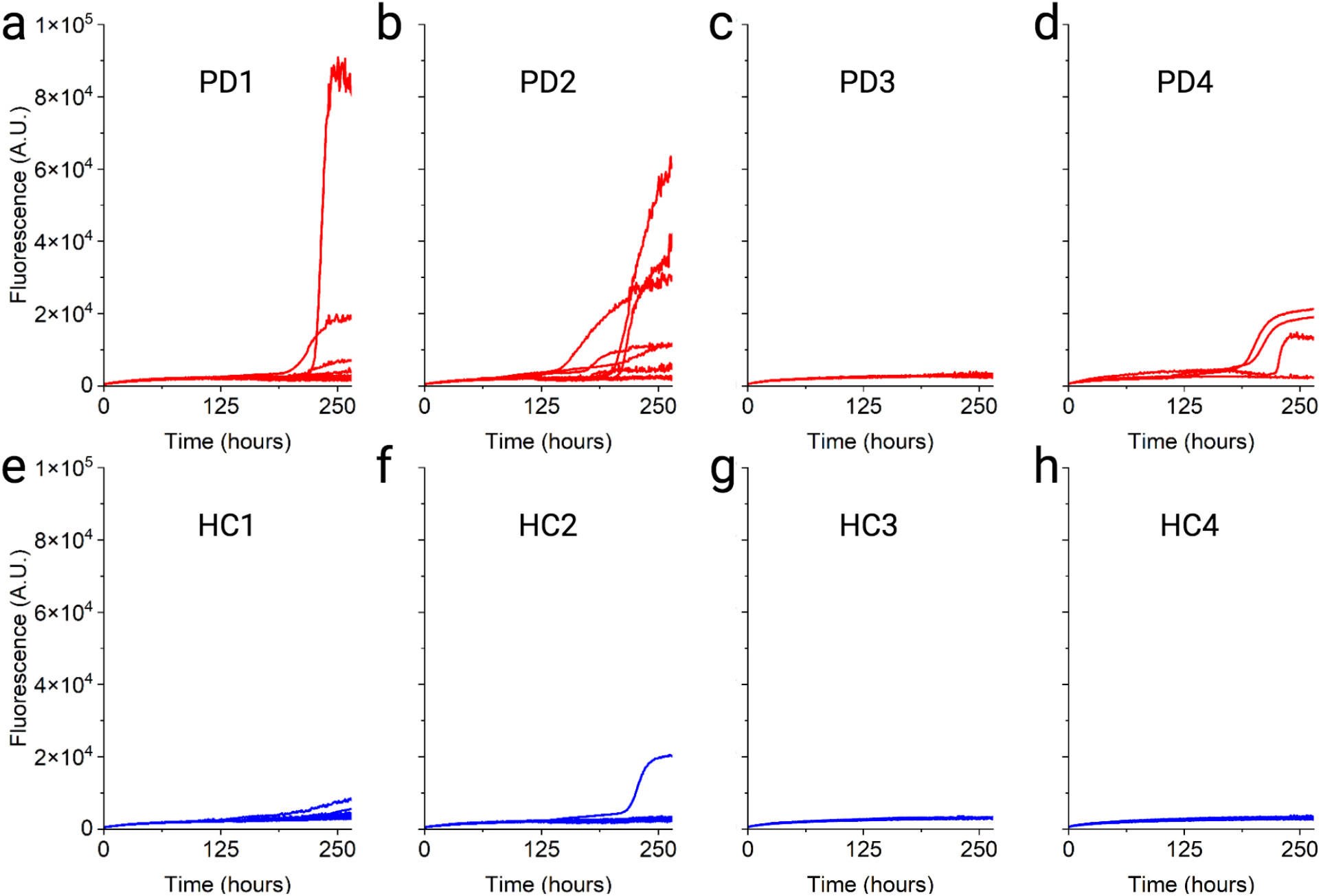
(a-d) Nano-QuIC fluorescent kinetic curves of plasma from patients diagnosed with Parkinson’s disease (e-h) Nano-QuIC fluorescent kinetic curves of plasma from healthy control patients. (a,b,e&f n=10; c,d,g&h n=4) Scales are equal for all panels.

All plasma samples were centrifuged and all but 20ul of the supernatant removed. These samples were then resuspended and added to wells containing recombinant α-syn and silica nanoparticles (see methods for details). Due to limited sample availability, HC3, HC4, PD3 & PD4 only had 4 technical replicates compared to 10 technical replicates for other samples. Utilizing the fluorescence curves (Fig. 5a-h), there was clearly increased seeding activity in three out of the four PD samples versus the four HC samples.

## Discussion

Our study shows the enhanced efficiency of Nano-QuIC reactions utilizing 50 nm siNPs, which significantly expedited the rate of amyloid formation of artificially misfolded seeds compared to reactions with no SiNPs (cutting times to detection in half) while displaying minimal false seeding. By sweeping the pH of our Nano-QuIC reactions, we showed that seeding activity is more efficiently detected at lower pH, supporting the hypothesis that electrostatic charge plays a role in the mechanism of Nano-QuIC. We successfully leveraged Nano-QuIC to detect misfolded α-syn as low as 900 pg/ml in human blood plasma (Fig. 3) and 90 pg/ml in lysed whole bovine blood (Fig. 4), demonstrating the potential use of Nano-QuIC for detection of disease associated α-syn in blood.

Human plasma, often used in routine diagnostic assays due to its noninvasive acquisition process, represents an ideal target for assay development. Whole bovine blood was used because it serves as an excellent model displaying the complexity of real biological samples. Lysed blood maximizes inhibitors, thus representing an extremely challenging sample. The abundance of bovine blood also facilitates the early stages of our protocol development.

We observed the 50nm siNPs in reactions at pH 6.3 gave the fastest kinetics while maintaining minimal false seeding (Fig. 2). We reasoned that the siNPs accelerated the reaction kinetics of misfolded protein aggregation because substrate proteins adsorbed onto the silica surface partly through electrostatic interactions.^37^ In previous work, we showed that α-syn can bind to the inner lumen of glass capillary tubes.^42^ On SiNPs, the adsorbed protein is more concentrated at the surface and likely changes conformation to a state more readily misfolded^43– 45^ (Fig. 1). Above the isoelectric point of α-syn (∼4.7), the net charge of α-syn is negative, which likely decreases the amount of α-syn interacting with the negatively charged siNPs. As the pH decreases, the net charge on the α-syn becomes closer to neutral, thus allowing it to interact more efficiently with the negatively charged siNPs, possibly through hydrophobic, hydrophilic, and local electrostatic interactions.^38,46^ This trend, reflected in Fig. 2b, shows that as the pH of the solution decreases the interactions of the α-syn with the siNPs increases, as seen by the increased rate of amyloid formation. Additionally, it has been reported that α-syn can interact with negatively charged gold nanoparticles at pH around 6.2. ^47^ The false seeding observed with the 20 nm SiNPs (Fig 2a) could be due to their curvature or perhaps the greater surface area available for proteins to interact with. Lynch et al. ^48^ note that curvature has a profound impact on protein secondary structure.

We have demonstrated that Nano-QuIC is a promising technique to overcome inhibitory barriers commonly associated with blood/plasma samples in conventional amplification assays. Whole blood samples are extremely complex and challenging for diagnostics. In samples such as CSF, blood contamination is thought to greatly interfere with QuIC diagnostics.^25,41^ Even in the presence robust inhibitors present in whole blood, Nano-QuIC successfully detected misfolded α-syn down to 90 pg/ml, whereas reactions without SiNPs failed to detect any presence at any dilutions (Fig. 4a-b). With human plasma, Nano-QuIC again outperformed reactions without SiNPs, having no false positive replicates while detecting misfolded α-syn at a level 900 pg/ml, a 100-fold lower than reactions containing no SiNPs (Fig. 3a-d). It should be noted that some synucleinopathy amplification assays utilize millimeter-scale beads ^12,21,22,49,50^ in the reaction while others do not ^20,51–54^. Additionally, RT-QuIC protocols for other protein misfolding diseases such as CJD, ALS, Alzheimer’s, and CWD have not utilized mm scale beads in reactions.^55–59^ In this study, beads were left out of the reaction to isolate the effect of nanoparticles on protein amplification assays. Future studies should examine the effects of utilizing mm scale beads and nanoparticles to produce a more optimized assay.

Finally we tested human plasma from patients diagnosed with PD versus healthy controls. By observing the fluorescent curves, it was seen that the PD samples showed higher seeding activity (Fig. 5). The seeding activity took between 150-230 hours to observe (Fig. 5a-d) which is longer than needed for the spiking plasma experiments in Fig. 3, however it is still clearly present. Longer seeding times are possibly in-part due to an extremely low quantity of misfolded α-syn in the plasma combined with a high concentration of inhibitors. Additionally for HC2, one well showed seeding activity. This is the only well that showed seeding out of the 28 technical replicates from the four HC samples. This single event, is likely random seeding, but could also be due to a low level of misfolded alpha-syn starting in that sample. One PD sample exhibited no seeding activity thus further optimization of temperature, rigor of shaking and SDS concentration needs to be done before clinical relevance. Nevertheless, these results serve as an early proof-of-concept experiment showing the potential of Nano-QuIC to detect misfolded alpha synuclein in plasma from patients with synucleinopathies.

In conclusion, the unique ability of Nano-QuIC to overcome inhibitors and accelerate detection time collectively mark a significant advance over traditional diagnostic methods without SiNPs. These improvements are especially useful for blood-based samples, which have historically been plagued by inhibitors that render them unsuitable for clinical applications of amplification assays such as PMCA or RT-QuIC. The promising results from our proof-of-concept experiments set the foundation for subsequent research.

More work needs to be done to develop this assay further. Firstly, the four PD and four HC samples in this work only show a proof-of-concept detection method in plasma; rigorous testing on a large cohort of patients must be performed to establish the sensitivity of this protocol. Assays requiring 270 hours, while not completely out of the question^20,54^, would be inconvenient for the clinical application; thus, it is important that future work focuses on speeding up the assay. Additionally, while other studies have found that amplification assays can distinguish between synucleinopathies and other protein misfolding disorders^21,51,54^, controls such as Alzheimer’s, CJD, ALS, etc., should be included to determine Nano-QuIC’s specificity. Further studies should then focus on Nano-QuIC’s ability to distinguish between synucleinopathy diseases such as PD, MSA, DLB, etc. If it is found that Nano-QuIC can distinguish between synucleinopathies, then it could be an extremely valuable tool to help clinicians determine disease. If it is not specific to any particular synucleinopathy, then it could be used as a screening tool to indicate if further testing is needed. Either way, with future development and rigorous validation we envision Nano-QuIC to be a valuable tool to aid the synucleinopathy field. Should future Nano-QuIC protocols become integrated into clinical diagnostics, it could not only facilitate early disease detection but also aid in patient prognosis, and assess drug efficacy, potentially impacting the way we could manage these debilitating conditions.

## Supporting Information

Methods section summarizing sample preparation, experimental Nano-QuIC protocols, and data analysis.

## Author Contributions

S.H.O. conceived the study. P.R.C., H.J, H.A. and M.L. performed molecular experiments. P.R.C., M.L., H.J, H.A., P.M., R.S., P.L., W.S., S.H.O., H.Y.P. and P.A.L. assisted with experimental design and interpreted the results. D.L. provided human blood plasma samples for spiking experiments. A.S. coordinated Parkinson’s patient sample logistics. P.R.C. and M.L. performed statistical analyses. S.H.O., W.S., H.Y.P. and P.A.L. oversaw the research. All authors wrote and contributed to the final manuscript.

## Notes

P.R.C., M.L., P.A.L., and S.-H.O. declare the following competing financial interest: A provisional patent has been filed, which covers specific ideas, design, and protocols outlined within this paper. All other authors declare no competing interests. P.A.L. and S.-H.O. are co-founders and equity holders of Priogen Corp, a diagnostic company specializing in the detection of prions and protein-misfolding diseases.

## Acknowledgments

The authors thank NIH Rocky Mountain Labs, especially Byron Caughey, Andrew Hughson, and Christina Orrù for training and assistance with the implementation of RT-QuIC. Luciano Caixeta provided bovine blood samples. We would like to thank the Micheal J Fox foundation for connecting us to the National Institute of Neurological Disorders and Stroke. We would like to thank the National Institute of Neurological Disorders and Stroke and the Indiana University School of Medicine for facilitating and providing pooled human plasma samples for initial protocol development. Gage Rowden provided valuable assistance. S. Stone provided valuable logistical assistance with our molecular work. Some figures were created with Biorender.com.

## References

(1) Lashuel, H. A.; Overk, C. R.; Oueslati, A.; Masliah, E. The Many Faces of α-Synuclein: From Structure and Toxicity to Therapeutic Target. Nat. Rev. Neurosci. 2013, 14 (1), 38– 48.

(2) Ebrahimi-Fakhari, D.; Wahlster, L.; McLean, P. J. Protein Degradation Pathways in Parkinson’s Disease: Curse or Blessing. Acta Neuropathol. 2012, 124 (2), 153–172.

(3) Srivastava, A.; Alam, P.; Caughey, B. RT-QuIC and Related Assays for Detecting and Quantifying Prion-like Pathological Seeds of α-Synuclein. Biomolecules 2022, 12 (4), 576.

(4) Experimental Models of Parkinson’s Disease, 1st ed.; Imai, Y., Ed.; Methods in molecular biology (Clifton, N.J.); Springer: New York, NY, 2021.

(5) Parnetti, L.; Gaetani, L.; Eusebi, P.; Paciotti, S.; Hansson, O.; El-Agnaf, O.; Mollenhauer, B.; Blennow, K.; Calabresi, P. CSF and Blood Biomarkers for Parkinson’s Disease. Lancet Neurol. 2019, 18 (6), 573–586.

(6) Siderowf, A.; Concha-Marambio, L.; Lafontant, D.-E.; Farris, C. M.; Ma, Y.; Urenia, P. A.; Nguyen, H.; Alcalay, R. N.; Chahine, L. M.; Foroud, T.; Galasko, D.; Kieburtz, K.; Merchant, K.; Mollenhauer, B.; Poston, K. L.; Seibyl, J.; Simuni, T.; Tanner, C. M.; Weintraub, D.; Videnovic, A.; Choi, S. H.; Kurth, R.; Caspell-Garcia, C.; Coffey, C. S.; Frasier, M.; Oliveira, L. M. A.; Hutten, S. J.; Sherer, T.; Marek, K.; Soto, C.; Parkinson’s Progression Markers Initiative. Assessment of Heterogeneity among Participants in the Parkinson’s Progression Markers Initiative Cohort Using α-Synuclein Seed Amplification: A Cross-Sectional Study. Lancet Neurol. 2023, 22 (5), 407–417.

(7) Simuni, T.; Chahine, L. M.; Poston, K.; Brumm, M.; Buracchio, T.; Campbell, M.; Chowdhury, S.; Coffey, C.; Concha-Marambio, L.; Dam, T.; DiBiaso, P.; Foroud, T.; Frasier, M.; Gochanour, C.; Jennings, D.; Kieburtz, K.; Kopil, C. M.; Merchant, K.; Mollenhauer, B.; Montine, T.; Nudelman, K.; Pagano, G.; Seibyl, J.; Sherer, T.; Singleton, A.; Stephenson, D.; Stern, M.; Soto, C.; Tanner, C. M.; Tolosa, E.; Weintraub, D.; Xiao, Y.; Siderowf, A.; Dunn, B.; Marek, K. A Biological Definition of Neuronal α-Synuclein Disease: Towards an Integrated Staging System for Research. Lancet Neurol. 2024, 23 (2), 178–190.

(8) Okuzumi, A.; Hatano, T.; Matsumoto, G.; Nojiri, S.; Ueno, S.-I.; Imamichi-Tatano, Y.; Kimura, H.; Kakuta, S.; Kondo, A.; Fukuhara, T.; Li, Y.; Funayama, M.; Saiki, S.; Taniguchi, D.; Tsunemi, T.; McIntyre, D.; Gérardy, J.-J.; Mittelbronn, M.; Kruger, R.; Uchiyama, Y.; Nukina, N.; Hattori, N. Propagative α-Synuclein Seeds as Serum Biomarkers for Synucleinopathies. Nat. Med. 2023, 29, 1448–1455.

(9) Delenclos, M.; Jones, D. R.; McLean, P. J.; Uitti, R. J. Biomarkers in Parkinson’s Disease: Advances and Strategies. Parkinsonism Relat. Disord. 2016, 22, S106–S110.

(10) Singer, W.; Schmeichel, A. M.; Shahnawaz, M.; Schmelzer, J. D.; Sletten, D. M.; Gehrking, T. L.; Gehrking, J. A.; Olson, A. D.; Suarez, M. D.; Misra, P. P.; Soto, C.; Low, P. A. Alpha-Synuclein Oligomers and Neurofilament Light Chain Predict Phenoconversion of Pure Autonomic Failure. Ann. Neurol. 2021, 89 (6), 1212–1220.

(11) Singer, W.; Schmeichel, A. M.; Shahnawaz, M.; Schmelzer, J. D.; Boeve, B. F.; Sletten, D. M.; Gehrking, T. L.; Gehrking, J. A.; Olson, A. D.; Savica, R.; Suarez, M. D.; Soto, C.; Low, P. A. Alpha-Synuclein Oligomers and Neurofilament Light Chain in Spinal Fluid Differentiate Multiple System Atrophy from Lewy Body Synucleinopathies. Ann. Neurol. 2020, 88 (3), 503–512.

(12) Concha-Marambio, L.; Pritzkow, S.; Shahnawaz, M.; Farris, C. M.; Soto, C. Seed Amplification Assay for the Detection of Pathologic Alpha-Synuclein Aggregates in Cerebrospinal Fluid. Nat. Protoc. 2023, 18 (4), 1179–1196.

(13) Lee, D. J.; Christenson, P. R.; Rowden, G.; Lindquist, N. C.; Larsen, P. A.; Oh, S.-H. Rapid on-Site Amplification and Visual Detection of Misfolded Proteins via Microfluidic Quaking-Induced Conversion (Micro-QuIC). npj Biosensing 2024, 1, 6.

(14) Candelise, N.; Schmitz, M.; Thüne, K.; Cramm, M.; Rabano, A.; Zafar, S.; Stoops, E.; Vanderstichele, H.; Villar-Pique, A.; Llorens, F.; Zerr, I. Effect of the Micro-Environment on α-Synuclein Conversion and Implication in Seeded Conversion Assays. Transl. Neurodegener. 2020, 9, 5.

(15) Candelise, N.; Baiardi, S.; Franceschini, A.; Rossi, M.; Parchi, P. Towards an Improved Early Diagnosis of Neurodegenerative Diseases: The Emerging Role of in Vitro Conversion Assays for Protein Amyloids. Acta Neuropathol Commun 2020, 8 (1), 117.

(16) Coughlin, D. G.; Irwin, D. J. Fluid and Biopsy Based Biomarkers in Parkinson’s Disease. Neurotherapeutics 2023, 20, 932–954.

(17) Bongianni, M.; Catalan, M.; Perra, D.; Fontana, E.; Janes, F.; Bertolotti, C.; Sacchetto, L.; Capaldi, S.; Tagliapietra, M.; Polverino, P.; Tommasini, V.; Bellavita, G.; Kachoie, E. A.; Baruca, R.; Bernardini, A.; Valente, M.; Fiorini, M.; Bronzato, E.; Tamburin, S.; Bertolasi, L.; Brozzetti, L.; Cecchini, M. P.; Gigli, G.; Monaco, S.; Manganotti, P.; Zanusso, G. Olfactory Swab Sampling Optimization for α-Synuclein Aggregate Detection in Patients with Parkinson’s Disease. Transl. Neurodegener. 2022, 11 (1), 37.

(18) Manne, S.; Kondru, N.; Jin, H.; Serrano, G. E.; Anantharam, V.; Kanthasamy, A.; Adler, C. H.; Beach, T. G.; Kanthasamy, A. G. Blinded RT-QuIC Analysis of α-Synuclein Biomarker in Skin Tissue From Parkinson’s Disease Patients. Mov. Disord. 2020, 35 (12), 2230–2239.

(19) De Luca, C. M. G.; Elia, A. E.; Portaleone, S. M.; Cazzaniga, F. A.; Rossi, M.; Bistaffa, E.; De Cecco, E.; Narkiewicz, J.; Salzano, G.; Carletta, O.; Romito, L.; Devigili, G.; Soliveri, P.; Tiraboschi, P.; Legname, G.; Tagliavini, F.; Eleopra, R.; Giaccone, G.; Moda, F. Efficient RT-QuIC Seeding Activity for α-Synuclein in Olfactory Mucosa Samples of Patients with Parkinson’s Disease and Multiple System Atrophy. Transl. Neurodegener. 2019, 8, 24.

(20) Shahnawaz, M.; Mukherjee, A.; Pritzkow, S.; Mendez, N.; Rabadia, P.; Liu, X.; Hu, B.; Schmeichel, A.; Singer, W.; Wu, G.; Tsai, A.-L.; Shirani, H.; Nilsson, K. P. R.; Low, P. A.; Soto, C. Discriminating α-Synuclein Strains in Parkinson’s Disease and Multiple System Atrophy. Nature 2020, 578 (7794), 273–277.

(21) Groveman, B. R.; Orrù, C. D.; Hughson, A. G.; Raymond, L. D.; Zanusso, G.; Ghetti, B.; Campbell, K. J.; Safar, J.; Galasko, D.; Caughey, B. Rapid and Ultra-Sensitive Quantitation of Disease-Associated α-Synuclein Seeds in Brain and Cerebrospinal Fluid by αSyn RT-QuIC. Acta Neuropathol Commun 2018, 6 (1), 7.

(22) Fairfoul, G.; McGuire, L. I.; Pal, S.; Ironside, J. W.; Neumann, J.; Christie, S.; Joachim, C.; Esiri, M.; Evetts, S. G.; Rolinski, M.; Baig, F.; Ruffmann, C.; Wade-Martins, R.; Hu, M. T. M.; Parkkinen, L.; Green, A. J. E. Alpha-Synuclein RT-QuIC in the CSF of Patients with Alpha-Synucleinopathies. Ann Clin Transl Neurol 2016, 3 (10), 812–818.

(23) Nakai, M.; Fujita, M.; Waragai, M.; Sugama, S.; Wei, J.; Akatsu, H.; Ohtaka-Maruyama, C.; Okado, H.; Hashimoto, M. Expression of Alpha-Synuclein, a Presynaptic Protein Implicated in Parkinson’s Disease, in Erythropoietic Lineage. Biochem. Biophys. Res. Commun. 2007, 358 (1), 104–110.

(24) Abd Elhadi, S.; Grigoletto, J.; Poli, M.; Arosio, P.; Arkadir, D.; Sharon, R. α-Synuclein in Blood Cells Differentiates Parkinson’s Disease from Healthy Controls. Ann Clin Transl Neurol 2019, 6 (12), 2426–2436.

(25) Cramm, M.; Schmitz, M.; Karch, A.; Mitrova, E.; Kuhn, F.; Schroeder, B.; Raeber, A.; Varges, D.; Kim, Y.-S.; Satoh, K.; Collins, S.; Zerr, I. Stability and Reproducibility Underscore Utility of RT-QuIC for Diagnosis of Creutzfeldt-Jakob Disease. Mol. Neurobiol. 2016, 53 (3), 1896–1904.

(26) Kuzkina, A.; Bargar, C.; Schmitt, D.; Rößle, J.; Wang, W.; Schubert, A.-L.; Tatsuoka, C.; Gunzler, S. A.; Zou, W.-Q.; Volkmann, J.; Sommer, C.; Doppler, K.; Chen, S. G. Diagnostic Value of Skin RT-QuIC in Parkinson’s Disease: A Two-Laboratory Study. NPJ Parkinsons Dis 2021, 7 (1), 99.

(27) Bargar, C.; Wang, W.; Gunzler, S. A.; LeFevre, A.; Wang, Z.; Lerner, A. J.; Singh, N.; Tatsuoka, C.; Appleby, B.; Zhu, X.; Xu, R.; Haroutunian, V.; Zou, W.-Q.; Ma, J.; Chen, S. G. Streamlined Alpha-Synuclein RT-QuIC Assay for Various Biospecimens in Parkinson’s Disease and Dementia with Lewy Bodies. Acta Neuropathol Commun 2021, 9 (1), 62.

(28) Altug, H.; Oh, S.-H.; Maier, S. A.; Homola, J. Advances and Applications of Nanophotonic Biosensors. Nat. Nanotechnol. 2022, 17 (1), 5–16.

(29) Henderson, D. M.; Davenport, K. A.; Haley, N. J.; Denkers, N. D.; Mathiason, C. K.; Hoover, E. A. Quantitative Assessment of Prion Infectivity in Tissues and Body Fluids by Real-Time Quaking-Induced Conversion. J. Gen. Virol. 2015, 96 (Pt 1), 210–219.

(30) Davenport, K. A.; Hoover, C. E.; Denkers, N. D.; Mathiason, C. K.; Hoover, E. A. Modified Protein Misfolding Cyclic Amplification Overcomes Real-Time Quaking-Induced Conversion Assay Inhibitors in Deer Saliva To Detect Chronic Wasting Disease Prions. J. Clin. Microbiol. 2018, 56 (9), e00947–18.

(31) Hoover, C. E.; Davenport, K. A.; Henderson, D. M.; Zabel, M. D.; Hoover, E. A. Endogenous Brain Lipids Inhibit Prion Amyloid Formation In Vitro. J. Virol. 2017, 91 (9), 1– 12.

(32) Li, M.; Schwabenlander, M. D.; Rowden, G. R.; Schefers, J. M.; Jennelle, C. S.; Carstensen, M.; Seelig, D.; Larsen, P. A. RT-QuIC Detection of CWD Prion Seeding Activity in White-Tailed Deer Muscle Tissues. Sci. Rep. 2021, 11 (1), 16759.

(33) Bistaffa, E.; Vuong, T. T.; Cazzaniga, F. A.; Tran, L.; Salzano, G.; Legname, G.; Giaccone, G.; Benestad, S. L.; Moda, F. Use of Different RT-QuIC Substrates for Detecting CWD Prions in the Brain of Norwegian Cervids. Sci. Rep. 2019, 9 (1), 18595.

(34) Kluge, A.; Bunk, J.; Schaeffer, E.; Drobny, A.; Xiang, W.; Knacke, H.; Bub, S.; Lückstädt, W.; Arnold, P.; Lucius, R.; Berg, D.; Zunke, F. Detection of Neuron-Derived Pathological α-Synuclein in Blood. Brain 2022, 145 (9), 3058–3071.

(35) Kluge, A.; Schaeffer, E.; Bunk, J.; Sommerauer, M.; Röttgen, S.; Schulte, C.; Roeben, B.; von Thaler, A.-K.; Welzel, J.; Lucius, R.; Heinzel, S.; Xiang, W.; Eschweiler, G. W.; Maetzler, W.; Suenkel, U.; Berg, D. Detecting Misfolded α-Synuclein in Blood Years before the Diagnosis of Parkinson’s Disease. Mov. Disord. 2024, 39, 1289–1299.

(36) Kumar, A.; Kim, S.; Nam, J.-M. Plasmonically Engineered Nanoprobes for Biomedical Applications. J. Am. Chem. Soc. 2016, 138 (44), 14509–14525.

(37) Christenson, P. R.; Li, M.; Rowden, G.; Larsen, P. A.; Oh, S.-H. Nanoparticle-Enhanced RT-QuIC (Nano-QuIC) Diagnostic Assay for Misfolded Proteins. Nano Lett. 2023, 23 (9), 4074–4081.

(38) Christenson, P. R.; Li, M.; Rowden, G.; Schwabenlander, M. D.; Wolf, T. M.; Oh, S.-H.; Larsen, P. A. A Field-Deployable Diagnostic Assay for the Visual Detection of Misfolded Prions. Sci. Rep. 2022, 12 (1), 12246.

(39) Buell, A. K.; Galvagnion, C.; Gaspar, R.; Sparr, E.; Vendruscolo, M.; Knowles, T. P. J.; Linse, S.; Dobson, C. M. Solution Conditions Determine the Relative Importance of Nucleation and Growth Processes in α-Synuclein Aggregation. Proc. Natl. Acad. Sci. U. S. A. 2014, 111 (21), 7671–7676.

(40) Coysh, T.; Mead, S. The Future of Seed Amplification Assays and Clinical Trials. Front. Aging Neurosci. 2022, 14, 872629.

(41) Dong, T.-T.-T.; Satoh, K. The Latest Research on RT-QuIC Assays-A Literature Review. Pathogens 2021, 10 (3), 305.

(42) Christenson, P. R.; Jeong, H.; Ahn, H.; Li, M.; Rowden, G.; Shoemaker, R. L.; Larsen, P. A.; Park, H. Y.; Oh, S.-H. Visual Detection of Misfolded Alpha-Synuclein and Prions via Capillary-Based Quaking-Induced Conversion Assay (Cap-QuIC). npj Biosensing 2024, 1, 2.

(43) Grigolato, F.; Arosio, P. The Role of Surfaces on Amyloid Formation. Biophys. Chem. 2021, 270, 106533.

(44) Zhang, D.; Neumann, O.; Wang, H.; Yuwono, V. M.; Barhoumi, A.; Perham, M.; Hartgerink, J. D.; Wittung-Stafshede, P.; Halas, N. J. Gold Nanoparticles Can Induce the Formation of Protein-Based Aggregates at Physiological pH. Nano Lett. 2009, 9 (2), 666–671.

(45) John, T.; Gladytz, A.; Kubeil, C.; Martin, L. L.; Risselada, H. J.; Abel, B. Impact of Nanoparticles on Amyloid Peptide and Protein Aggregation: A Review with a Focus on Gold Nanoparticles. Nanoscale 2018, 10 (45), 20894–20913.

(46) Kim, Y.; Park, J.-H.; Lee, H.; Nam, J.-M. How Do the Size, Charge and Shape of Nanoparticles Affect Amyloid β Aggregation on Brain Lipid Bilayer? Sci. Rep. 2016, 6, 19548.

(47) Alvarez, Y. D.; Fauerbach, J. A.; Pellegrotti, J. V.; Jovin, T. M.; Jares-Erijman, E. A.; Stefani, F. D. Influence of Gold Nanoparticles on the Kinetics of α-Synuclein Aggregation. Nano Lett. 2013, 13 (12), 6156–6163.

(48) Lynch, I.; Dawson, K. A. Protein–Nanoparticle Interactions. In Nano-Enabled Medical Applications; Jenny Stanford Publishing, 2020; pp 231–250.

(49) Hall, S.; Orrù, C. D.; Serrano, G. E.; Galasko, D.; Hughson, A. G.; Groveman, B. R.; Adler, C. H.; Beach, T. G.; Caughey, B.; Hansson, O. Performance of αSynuclein RT-QuIC in Relation to Neuropathological Staging of Lewy Body Disease. Acta Neuropathol. Commun. 2022, 10 (1), 90.

(50) Peña-Bautista, C.; Kumar, R.; Baquero, M.; Johansson, J.; Cháfer-Pericás, C.; Abelein, A.; Ferreira, D. Misfolded Alpha-Synuclein Detection by RT-QuIC in Dementia with Lewy Bodies: A Systematic Review and Meta-Analysis. Front. Mol. Biosci. 2023, 10, 1193458.

(51) Wang, Z.; Becker, K.; Donadio, V.; Siedlak, S.; Yuan, J.; Rezaee, M.; Incensi, A.; Kuzkina, A.; Orrú, C. D.; Tatsuoka, C.; Liguori, R.; Gunzler, S. A.; Caughey, B.; Jimenez-Capdeville, M. E.; Zhu, X.; Doppler, K.; Cui, L.; Chen, S. G.; Ma, J.; Zou, W.-Q. Skin α-Synuclein Aggregation Seeding Activity as a Novel Biomarker for Parkinson Disease. JAMA Neurol. 2021, 78 (1), 30–40.

(52) Sano, K.; Atarashi, R.; Satoh, K.; Ishibashi, D.; Nakagaki, T.; Iwasaki, Y.; Yoshida, M.; Murayama, S.; Mishima, K.; Nishida, N. Prion-like Seeding of Misfolded α-Synuclein in the Brains of Dementia with Lewy Body Patients in RT-QUIC. Mol. Neurobiol. 2018, 55 (5), 3916–3930.

(53) Kuang, Y.; Mao, H.; Gan, T.; Guo, W.; Dai, W.; Huang, W.; Wu, Z.; Li, H.; Huang, X.; Yang, X.; Xu, P.-Y. A Skin-Specific α-Synuclein Seeding Amplification Assay for Diagnosing Parkinson’s Disease. npj Parkinson’s Disease 2024, 10, 129.

(54) Shahnawaz, M.; Tokuda, T.; Waragai, M.; Mendez, N.; Ishii, R.; Trenkwalder, C.; Mollenhauer, B.; Soto, C. Development of a Biochemical Diagnosis of Parkinson Disease by Detection of α-Synuclein Misfolded Aggregates in Cerebrospinal Fluid. JAMA Neurol. 2017, 74 (2), 163–172.

(55) Mielke, J. K.; Klingeborn, M.; Schultz, E. P.; Markham, E. L.; Reese, E. D.; Alam, P.; Mackenzie, I. R.; Ly, C. V.; Caughey, B.; Cashman, N. R.; Leavens, M. J. Seeding Activity of Human Superoxide Dismutase 1 Aggregates in Familial and Sporadic Amyotrophic Lateral Sclerosis Postmortem Neural Tissues by Real-Time Quaking-Induced Conversion. Acta Neuropathol. 2024, 147 (1), 100.

(56) Raymond, G. J.; Race, B.; Orrú, C. D.; Raymond, L. D.; Bongianni, M.; Fiorini, M.; Groveman, B. R.; Ferrari, S.; Sacchetto, L.; Hughson, A. G.; Monaco, S.; Pocchiari, M.; Zanusso, G.; Caughey, B. Transmission of CJD from Nasal Brushings but Not Spinal Fluid or RT-QuIC Product. Ann Clin Transl Neurol 2020, 7 (6), 932–944.

(57) Tennant, J. M.; Henderson, D. M.; Wisniewski, T. M.; Hoover, E. A. RT-QuIC Detection of Tauopathies Using Full-Length Tau Substrates. Prion 2020, 14 (1), 249–256.

(58) Cazzaniga, F. A.; Bistaffa, E.; De Luca, C. M. G.; Bufano, G.; Indaco, A.; Giaccone, G.; Moda, F. Sporadic Creutzfeldt-Jakob Disease: Real-Time Quaking Induced Conversion (RT-QuIC) Assay Represents a Major Diagnostic Advance. Eur. J. Histochem. 2021, 65 (1), 3298.

(59) Thomas, C. M.; Salamat, M. K. F.; de Wolf, C.; McCutcheon, S.; Blanco, A. R. A.; Manson, J. C.; Hunter, N.; Houston, E. F. Development of a Sensitive Real-Time Quaking-Induced Conversion (RT-QuIC) Assay for Application in Prion-Infected Blood. PLoS One 2023, 18 (11), e0293845.

