## Supporting Information for "Blood-based Nano-QuIC: Inhibitor-resistant Detection of Seeding Activity in Patients Diagnosed with Parkinson’s Disease"

**Supporting Figure**

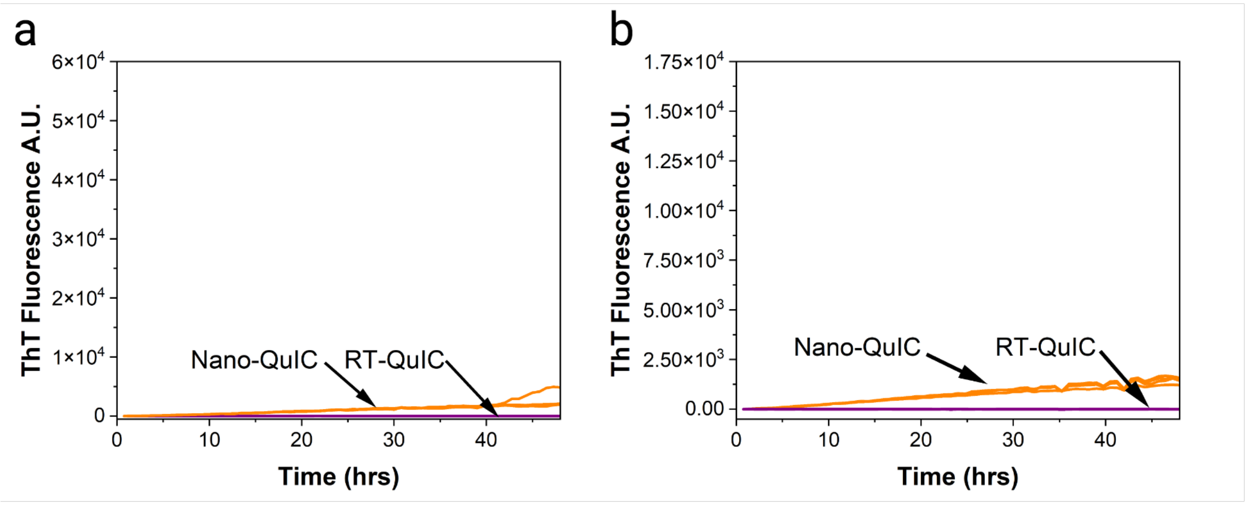


**Supporting Figure 1:** (a) Plasma artificial spiking experiment negative control curves for Nano-QuIC (n=5) and RT-QuIC (n=5). (b) Whole blood spiking experiment negative control curves for Nano-QuIC (n=4) and RT-QuIC (n=4). Note: scales for panel a & b match corresponding main figures 3 & 4 respectively.

### Methods

#### Preparation of spontaneously misfolded seed

To prepare misfolded α-syn for nanoparticle diameter, pH and plasma/blood spike experiments, a master mix was made to the following specifications: 1X Phosphate buffer (9.2mM $NaH_2PO_4$ , 2.8mM $Na_2HPO_4$ , 2.7mM KCl, 137mM NaCl), 1 mM Ethylenediaminetetraacetic acid (EDTA), 170 mM NaCl (in addition to that in the NaCl in the Phosphate buffer), 10 μM thioflavin T (ThT), 0.09 mg/mL recombinant human alpha synuclein (R&D systems). This was then shaken and incubated until spontaneous misfolding occurred. Wells were then pooled and 10 μL was added to 90 μL SDS/PBS (0.1% Sodium Dodecyl Sulfate [SDS], 1X PBS, 1X N-2 supplement (μL of N2 per 100 μL SDS/PBS) [Thermo Fisher Scientific, Waltham, MA, USA]). These samples were then shaken for 48 hrs at 48 C on a plate reader (BMG Labtech, Cary, North Carolina, USA; 700 rpm, double orbital, shake for 60 s, rest for 60 s). After this, samples were pooled and aliquoted for future use.

#### QuIC Size and pH

For Nano-QuIC analysis, a master mix was made to the following specifications: 1X Phosphate buffer (9.2mM $NaH_2PO_4$, 2.8mM $Na_2HPO_4$ , 2.7mM KCl, 137mM NaCl), 1 mM Ethylenediaminetetraacetic acid (EDTA), 170 mM NaCl (in addition to that in the NaCl in the Phosphate buffer), 10 μM thioflavin T (ThT), 0.09 mg/mL recombinant human alpha synuclein (R&D systems). For pH experiments the ratios of $NaH_2PO_4$ and $Na_2HPO_4$ were varied to give the range of pHs. pH was measured for solutions with the nanoparticles spun out. For size and concentration sweeps, silica nanoparticles from Fortis Life Sciences Company, San Diego, CA, USA with diameters ranging from 20nm-100nm were added to the reaction with final well concentrations of 2.5mg/mL, 0.5mg/mL, 0.1mg/mL or no NPs. For RT-QuIC reactions, the master mix was the same except with no NPs. 98 μL of the master mixes were added to wells on a 96 well plate. These wells were then spiked with 2 μL of seed solution (10 μL spontaneously misfolded human alpha syn in 90 μL of SDS/PBS).

**Human Plasma and bovine blood QuIC**

For human plasma experiments, EDTA plasma samples were obtained from Solomon Park Research Laboratories (Burien, WA, USA). Master mixes were prepared as described above to obtain a pH around 6.3. 10 µL of spontaneously misfolded α-syn (.09 mg/ml) was sonicated (200 W 15 sec on 15 sec off 3X times) and spiked into 90 µL of human plasma, this solution is referred to as $10^0$ dilution (Fig 4a). It should be noted that both spiking experiments were done with aliquots of the same spontaneously misfolded seed. The $10^0$ solution was then serially diluted tenfold in human plasma down to a final spike concentration of 90 pg/ml (referred to as $10^{-5}$). Each spike dilution was then diluted one hundred-fold in SDS/PBS (see above). 2 µL of this final dilution was then added to 98 µL of the master mix in a 96 well plate. Plates were put onto a plate reader (BMG Labtech, Cary, North Carolina, USA; 700 rpm, double orbital, shake for 60 s, rest for 60 s, gain 1000) at 42C for at least 48hrs.

For bovine whole blood dilutions, the protocol was the same except whole bovine blood lysed via freeze thaw cycles was used instead of human plasma.

**Human Plasma Parkinson's samples**

For human plasma sample testing, 240 ul of plasma was centrifuged at 21,000g for 40 minutes. 220 ul of supernatant was removed and the remaining liquid/pellet was resuspended in 20ul 0.1% SDS in 1X PBS. Master mixes were prepared using 1X phosphate buffer pH5.8 (9.2mM $NaH_2PO_4$, 2.8mM $Na_2HPO_4$ , 2.7mM KCl, 137mM NaCl), 1 mM Ethylenediaminetetraacetic acid (EDTA), 170 mM NaCl (in addition to that in the NaCl in the Phosphate buffer), 10 µM ThT, 0.1 mg/mL recombinant human alpha synuclein (R&D systems) and 2.5mg/ml 50 nm SiNPs. 2ul of resuspended plasma were added to wells containing mastermixes followed by the addition of 1ul of 0.1% SDS. For samples in main figure 6 a,b,e,f there were 10 technical replicates and for main figure 6 c,d,g,h there were 4 technical replicates, due to limited sample availability. Plates were put onto a plate reader (BMG Labtech, Cary, North Carolina, USA; 700 rpm, double orbital, shake for 60 s, rest for 60 s) at 42C for 265hrs.

**Evaluation of Kinetics Data**

Rate of amyloid formation was calculated as the inverse of the time to detection in seconds. The time to detection was classified as the start of the growth phase. The start of the growth phase was classified when the rate of change (rc) in fluorescence met the following conditions 1.) the rc is at least 100 units and the next data point has an rc of at least 250 units and 2.) the maximum rc was greater than 500 units. These conditions were used to take into account the differences in kinetic curve shapes (sharp, gentle slope…etc) of positive samples between experiments with pure artificial spiked seeds, lysed whole blood and plasma samples such that all experiments in the main manuscript could be analyzed with the same criteria.
